# In Silico Vaccine Design: Targeting Highly Epitopic Regions of *Clostridium perfringens* Type D Epsilon Toxin and *Clostridium novyi* Type B Alpha Toxin for Optimal Immunogenicity

**DOI:** 10.1101/2024.04.12.589226

**Authors:** Nastaran Ashoori, Mohammad Mehdi Ranjbar, Romana Schirhagl

**Author notes:** Corresponding Author: Nastaran Ashoori, Groningen University, University Medical Centre Groningen, Antonius Deusinglaan 1, 9713AW Groningen, the Netherlands.

## Abstract

Livestock infections caused by highly toxic bacteria pose significant challenges in veterinary medicine, often requiring complex and elusive treatment regimens. Developing effective vaccines tailored to combat these specific pathogens remains a pressing need within the field. Among the most formidable culprits are *Clostridium perfringens* type D and *Clostridium novyi* type B, notorious for their extreme toxicity and the difficulty in culturing them for vaccine production. In response to this challenge, our study endeavors to engineer a vaccine candidate capable of concurrently neutralizing the virulence of both bacterial strains. Leveraging computational techniques, we meticulously identified highly epitopic regions within *C. perfringens* Epsilon Toxin (ETX) and *C. novyi* Alpha Toxin (ATX), crucial targets for effective immunization. Through innovative fusion gene design, we integrated these epitopic regions alongside the PADRE-peptide sequence, serving as a universal adjuvant to bolster immune response. The culmination of our efforts materialized in the creation of Recombinant Fusion Protein D (rFPD), a novel vaccine construct poised to elicit robust and specific immune defenses against both bacterial species. By harnessing the power of in silico design and molecular engineering, our study heralds a promising stride towards mitigating the deleterious impact of livestock infections caused by these formidable pathogens.

## 1. Introduction

*Clostridium perfringens*, a Gram-positive, anaerobic, spore-forming, and non-motile bacterium, causes enteric diseases in domestic animals (Smedley J. G. and Fisher, 2005). Various strains of *C. perfringens* are categorized into five types (A to E) based on the toxins they produce. Recently, two additional types (F and G) have been identified for *C. perfringens* (Kaushik et al., 2019a; Rood et al., 2018). *C. perfringens* type D, the primary causative agent for enterotoxaemia in domestic ruminants, produces a 33 k D protoxin (328 aa residues). This protoxin undergoes cleavage and conversion into an active Epsilon toxin (ETX) by different proteases present in the host gastrointestinal tract, such as trypsin, chymotrypsin, or λ-protease encoded by *C. perfringens* (Rumah et al., 2013a). The pore-forming ETX considered the most toxic and fatal clostridial toxin after Botulinum and tetanus toxins, is a crucial virulence factor for *C. perfringens* disease in sheep and goats (Cole et al., 2004; Garcia et al., 2013; Kaushik et al., 2019a). The anaerobic environment in the gut of infected animals, attributed to the overfeeding of domestic ruminants, can lead to the overgrowth of *C. perfringens* type D (Kaushik et al., 2013) resulting in the excessive production of ETX. The gut mucosa absorbs ETX, leading to increased vascular permeability, elevated blood pressure, and intensive vascular damage, causing lesions in vital organs such as the heart, brain, kidneys, and lungs and the overproduction of ETX (Kaushik et al., 2019b; Miyamoto et al., 2008; Petit et al., 2003; Popoff, 2011; Tamai et al., 2003).

*Clostridium novyi* is the causative organism for gangrene in humans and animals (Gord-Noshahri et al., 2016). Based on the production of different toxins, *C. novyi* is categorized into 4 types (A-D) (Oakley et al., 1947, 1959; Willis & others, 1964). Types A, B and D are pathogenic, and type C is generally recognized as non-pathogenic (Nakamura et al., 1983; L. D. Smith et al., 1969; L. D. S. Smith & Hobbs, 1974). *C. novyi* type B, a Gram-positive, obligatory anaerobic, endospore-forming bacterium, causes necrotic hepatitis or black disease in domestic animals such as sheep, horses, pigs and cattle. The main virulence factor in *C. novyi* type B is Alpha toxin (ATX) (Hatheway, 1990). Production of ATX in the liver, after spore germination and development of the vegetative state, leads to necrotic infarction with wide regions of hyperemia (Felix et al., 2019; Kaushik et al., 2019b; Rumah et al., 2013b; Smedley J. G.and Fisher, 2005). ATX, a 200–250 kDa protein (2178 aa residues) belongs to the large clostridial cytotoxin family (Hofmann et al., 1995) and acts as a glycosyltransferase on Rho proteins in target cells. It disintegrates the cytoskeleton through inhibiting signal pathways resulting in vascular permeability and cell death (Amimoto et al., 1998; Bette et al., 1991; Gord-Noshahri et al., 2016; Hofmann et al., 1995).

In both ETX and ATX infections, the progression of the diseases is rapid, leading to animal death shortly after contamination. Therefore, using antibiotics is ineffective in treatment, and vaccination of domestic ruminants emerges as the most effective remedy (Felix et al., 2019; Kaushik et al., 2019b). Immunization with formaldehyde-inactivated toxins or toxoid-based vaccines, the current solution against *C. perfringens* and *C. novyi*, fails to induce the desired immunity against these bacteria. In other words, toxoids expose both immunoprotective and non-immunoprotective antigens to the body simultaneously, which is not only energy-consuming for the immune system but also triggers non-specific responses. Furthermore, improper inactivation of toxoids can lead to side effects. Given that both bacteria are anaerobes, cultivation and preparation of toxins become laborious and expensive (Aquino et al., 2016; Chandran et al., 2010; Gord-Noshahri et al., 2016; Kaushik et al., 2019b; Petit et al., 2003).

In recent years, recombinant DNA technology approaches have been employed for the development of a new generation of vaccines (Kaushik et al., 2013). Using this technique, recombinant Epsilon Toxin (rETX) has been produced and evaluated as a potential vaccine (Bokori-Brown et al., 2014). The study observed local inflammation, emphasizing the need for precise dosage determination to avoid potential damages. To address this challenge, the suggestion is to produce a small, highly immunogenic fragment of the toxin (Kaushik et al., 2019b).

Bioinformatics tools have been employed to design epitope-based vaccines, showing promise in comparison to conventional vaccines due to their safety, stability, and ease of production. Additionally, the accurate discovery of disease agents and critical epitopes facilitates specific immune responses against the organism, thereby preventing autoimmune responses and allergies (Hajighahramani et al., 2017; Mahmoodi et al., 2016; Negahdaripour et al., 2017, 2018; Nezafat et al., 2017; Soria-Guerra et al., 2015). In-silico approaches, instead of the time-consuming and challenging experimental methods, have been recently utilized to confidently identify accurate epitopes and provide faster outputs without incurring the risks and costs associated with culturing extremely toxic bacteria (Kaushik et al., 2019b; Negahdaripour et al., 2017).

The rational genetic combination of several genes coding for different proteins enables the production of polyvalent fusion proteins, such as multi-epitopic vaccines. Designing multi-target and highly efficient vaccines offers several benefits, including the accumulation of immunogenic B-cell and T-cell epitopes conserved in different variants of the pathogen within a single molecule. This approach allows for the removal of non-protective but immune-dominant epitopes, the addition of adjuvants to enhance immunogenicity, the use of linkers to join different segments together, thus avoiding the production of junctional epitopes, and an increase in the antigen presentation process (Negahdaripour et al., 2017; Nezafat et al., 2017).

In the present study, we designed and produced a fusion protein from epitopic regions of ETX and ATX, aiming to generate a potential vaccine against both *C. perfringens* and *C. novyi* infections simultaneously. We employed an effective rational design strategy by selecting and combining highly immunogenic regions using immunoinformatic and structural bioinformatics approaches. Specifically, kauwe designed a novel fusion gene, *rfpd*, encompassing epitopic regions of *C. perfringens* ETX, *C. novyi* ATX, and the PADRE-peptide sequence. Consequently, *rfpd* encodes the rFPD protein, which produces highly immunogenic regions based on B-cell epitopes of both toxins. T-helper epitopes are inserted into the construct to improve the vaccine immunogenicity of the two selected epitopic segments, along with the inclusion of PADRE-peptide as an adjuvant.

## 2. Materials and methods

We used immunoinformatic analysis to design a potential vaccine candidate against *C. perfringens* ETX and *C. novyi* ATX in silico. A schematic overview of the vaccine design process is presented in Figure 1.

**Figure 1.**
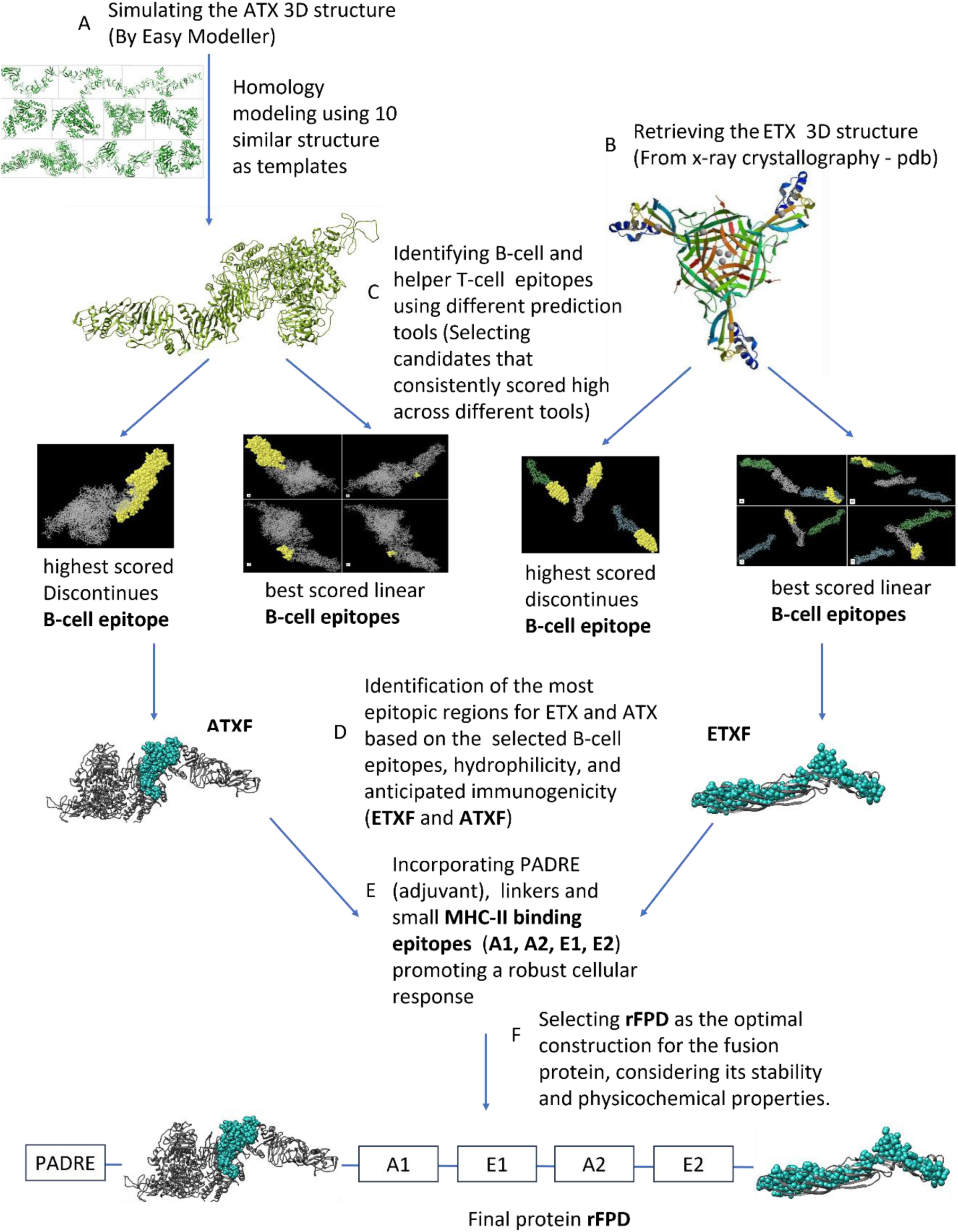
illustrates the in-silico vaccine design process against *C. perfringens* ETX and *C. novyi* ATX. The steps include A: Predicting and simulating the 3D structure of ATX using Easy Modeller. B: Retrieving the 3D structure (pdb) of ETX from the Protein Data Bank. C: Identifying different B-cell and helper T-cell epitopes of ETX and ATX using various prediction tools. D: Selecting the most epitopic regions of ETX and ATX based on chosen B-cell epitopes, hydrophilicity, and expected immunogenicity (ETXF and ATXF). E: Enhancing the construction by adding PADRE (adjuvant), linkers, and small MHC-II binding epitopes to activate a robust cellular response (A1, A2, E1, E2). F: Choosing rFPD as the best construction for the fusion protein based on its stability and physicochemical properties.

### 2.1 Primary analysis of ETX and ATX sequences

The nucleotide and amino acid sequences of both toxins were retrieved from the NCBI database and subjected to analysis using NCBI-Blast. For *C. perfringens* ETX, the NCBI reference sequence YP_002291114.1 and accession number KU726256.1 were utilized. The accession numbers Z48636.1 and CAA88565.1 were employed for *C. novyi* ATX. The secondary structure of both toxins, including β-sheet and α-helix formations, as well as their hydrophobicity (determined by the Kyte-Doolittle algorithm) and antigenicity (predicted using the Kolaskar-Tongaonkar algorithm), were assessed using the CLC Main Workbench 5.5. Areas of the proteins that are external and exposed, primarily hydrophilic, were identified as having high potential immunogenicity as B-cell epitopes.

### 2.2 Identification of the MHC-II binding epitopes

To predict helper T-cell binding MHC-II epitopes, the Immune Epitope Database (IEDB) with its MHC- II Binding Predictions server, TepiTool databases, and the CBS Prediction server (NetMHCII 2.2) were employed. The choice of H2-IAd, the most frequent allele in BALB/C mice (Ru, 2012), was made considering BALB/C mice as an animal model for injection and further experimental analysis. Among the available databases, IEDB was selected as the most comprehensive server for epitopic prediction algorithms (Mehla & Ramana, 2017). Consequently, the results from IEDB are considered valid and reliable. However, to ensure the selection of appropriate epitopes and validate the IEDB results, an assessment of the outcomes from various servers was undertaken.

### 2.3 Linear B-cell epitope prediction

For the prediction of linear epitopes of B-cell lymphocytes, the protein sequences of both toxins were submitted to various servers, including LBTope, SVMTriP, CBS-Bepipred, ABCpred, IEDB-Bepipred, IEDB-Kolaskar, and IEDB-Emini surface accessibility. These servers utilize the physicochemical properties of amino acid residues and machine learning algorithms to predict linear B-cell epitopes.

### 2.4 Determination of 3D Structures for Conformational B-Cell Epitope Prediction

To predict the conformational B-cell epitopes of ETX and ATX, the 3D structures of the toxins were subjected to relevant databases. The complete 3D model of *C. perfringens* ETX was retrieved from the Protein Data Bank (PDB) with PDB ID: 1UYJ. In the case of *C. novyi* ATX, where the 3D structure has not been obtained through crystallographic methods and lacks related reports, EasyModeller 4.0 software (Kuntal et al., 2010) was utilized for generating the 3D model based on homology modelling and multiple threading alignments. This software constructs 3D structures using the structural information of similar proteins in existing databases.

To find suitable toxin-like PDB structures for ATX modelling, the ATX amino acid sequence underwent NCBI-BLASTP 2.6.1, specifically limited to the Protein Data Bank server. From the blast result, 10 proteins were selected. Considering the amino acid sequence similarity of the chosen models with ATX exceeding 30%, the homology modelling method was deemed appropriate for generating a 3D model. The Swiss-Pdb Viewer 4.1.0 software was employed for energy-minimizing the predicted model. Subsequently, a Ramachandran plot analysis was conducted using RAMEPAGE to assess the accuracy of the 3D structure (Lovell et al., 2003). The validated model was then utilized for conformational B-cell epitope prediction.

### 2.5 Conformational B-cell epitope identification

The PDB file (3D structure) of ETX (PDB ID: 1UYJ) extracted from the PDB database, along with the final models of ATX, were submitted to CBTope, DiscoTope-2.0-CBS, and ElliPro servers for predicting conformational B-cell epitopes. CBTope employs the Support Vector Machine (SVM) method, utilizing amino acid composition calculations and recognizing antibody-interacting residues (Ansari & Raghava, 2010). DiscoTope servers function based on surface accessibility and propensity scores of each residue in the spatial vicinity (Kringelum et al., 2012). ElliPro predicts discontinuous epitopes by considering the accessibility and flexibility of residues in solution, providing a Protrusion Index (PI) score (Tolooe Zarin & Ahmadi, 2016).

Following the analysis of B-cell epitope predictions, one highly antigenic fragment from each toxin, exhibiting overlap in both continuous and discontinuous predicted epitopes, was selected as the most immunogenic part of the toxins. Antigenicity and hydrophobicity graphs, along with the secondary structure of the selected fragments of ETX and ATX, were compared with the main toxins.

### 2.6 The design of the recombinant fusion protein

The final selected B-cell epitopic fragments, along with the two best immunogenic helper T-cell epitopes (MHC-II Binding Epitopes) for each toxin, were considered. Appropriate linkers were chosen to fuse the selected segments. These linkers, acting as spacers, aid in the proper folding of the conformational epitopes in the chimeric proteins. To connect the MHC-II binding epitopes, the NH2- GPGPG-COOH linker was utilized, and for the B-cell epitopic fragments, the NH2-GGSSGG-COOH sequence was employed (Negahdaripour et al., 2017; Nezafat et al., 2017; Ranjbar et al., 2015).

The organization of the helper T-cell epitopes in the fusion protein was determined based on their scores. This decision was made considering that a stronger epitope, when positioned next to a weaker one, can attract immunodominance. To heighten the immunogenicity of the recombinant fusion protein, the PADRE sequence (universal T-helper Pan DR Epitope: AKFVAAWTLKAAA) was incorporated at the beginning of the structure. This tag can strongly and non-specifically bind to a variety of MHC-II alleles, reducing the risk of vaccine failure and acting as an adjuvant (Amimoto et al., 1998; Ghaffari-Nazari et al., 2015).

Five different models (Recombinant Fusion Protein A-D: rFPA, rFPB, rFPC, rFPD, and rFPE) were assessed for fusing the epitopes of both toxins to create the recombinant fusion protein. Their immunogenicity and physicochemical properties, including molecular weight, pH, half-life, solubility, and stability, were determined using the EXPASY-ProtParam tool. Additionally, protein allergenicity was examined through AlgPred and AllergenFP v.1.0 databases since the chosen model is not expected to be allergenic. The best model (rFPD) was selected based on its structure and a comprehensive consideration of all physicochemical properties.

### 2.7 Reverse translation and sequence verification of rFPD

The reverse translation of the rFPD structure into a nucleotide sequence was conducted. The nucleotide sequences of the selected components of the MHC-II binding epitopes, along with the B-cell epitopic regions, were extracted from the NCBI sequences for ETX (Accession Number: KU726256.1) and ATX (Accession Number: Z48636.1). However, the linkers and the PADRE peptide were reverse-translated using the BackTranseq-EMBOSS database. Subsequently, the JCat server was employed for codon usage adaptation, CG content optimization, and limiting restriction enzymes. Methionine nucleotide codon (ATG) as the start codon and the stop codon (TAG) were added to the constructs.

To examine the presence of any restriction enzyme sites in the gene, the final sequence was evaluated using CLC Main Workbench 5.5. This step is crucial, as the identification of restriction enzyme sites influences the selection of the most appropriate enzymes for cloning the gene into the vector. Additionally, to validate the correctness of the final sequence, this nucleotide sequence was translated into protein using CLC Main Workbench 5.5. The resulting protein sequence was then compared with our designed protein through BLAST in EXPASY-MultAlin.

## 3. Results

### 3.1. Primary analysis of the ETX and ATX sequences

The DNA and protein sequences of ETX and ATX, as analyzed through NCBI-Blast, demonstrate high conservation. This suggests that vaccine candidates utilizing epitopes from both toxins may be applicable against various strains of *C. perfringens* type D and *C. novyi* type B. For ETX, the hydrophobicity plot generated by CLC Main Workbench 5.5 indicates that, excluding the signal peptide (residues 1-32), the remaining protein is generally hydrophilic. The immunogenicity graph for ETX highlights the region around residues 150-300 as the most probable immunogenic area. The secondary structure analysis reveals that the residues after 135 aa predominantly form loops and β- sheets. Therefore, the region between 150-300 aa is considered the most epitopic area.

In the case of ATX, the hydrophobicity plot from CLC Main Workbench 5.5 indicates that the most hydrophilic area is approximately between residues 1200 and 2178. Despite different potential antigenic regions suggested by the antigenicity plot, we focus on epitopes related to the second half of the toxin, as it is more hydrophilic. The secondary structure prediction for ATX suggests that residues 1069-2178 mainly consist of β-sheets and loops. This region is considered promising for containing highly antigenic epitopes, given that loops and β-sheet structures are generally more antigenic than α-helix structures (Ingale, 2010).

### 3.2. Identification of MHC-II binding epitopes

MHC-II binding epitopes were predicted utilizing IEDB, TepiTool, and the CBS Prediction server for the prevalent MHC allele in mice (H2-IAd). Each server assigns numerical scores to the epitopes, where lower scores indicate stronger epitopes. The outcomes from various servers for both ETX and ATX were compared, and the two most highly immunogenic epitopes for each toxin were ultimately selected based on the IEDB results, considered the most reliable server (Supplementary material Appendix 1, Table S1).

### 3.3. Linear B-cell epitope prediction

Various servers (LBTope, SVMTriP, CBS-Bepipred, ABCpred, IEDB-Bepipred, IEDB-Kloskar, and IEDB- Emini surface accessibility) were employed to predict linear B-cell epitopes of ETX and ATX, with a threshold value set at 0.6. demonstrate that residues 150-300 aa for ETX and the C-terminal region between 1200-2178 aa in ATX encompass more continuous B-cell epitopes (Supplementary material Appendix 1, Table S2 and Table S3).

### 3.4. Determination of 3D structures for conformational B-cell epitope prediction

To predict discontinuous B-cell epitopes, servers require the 3D structure of the antigenic protein. The Protein Data Bank (PDB) provided the 3D structure for ETX (ETX-PDB) with PDB ID: 1UYJ (Cole et al., 2004) (Figure 2).

**Figure 2.**
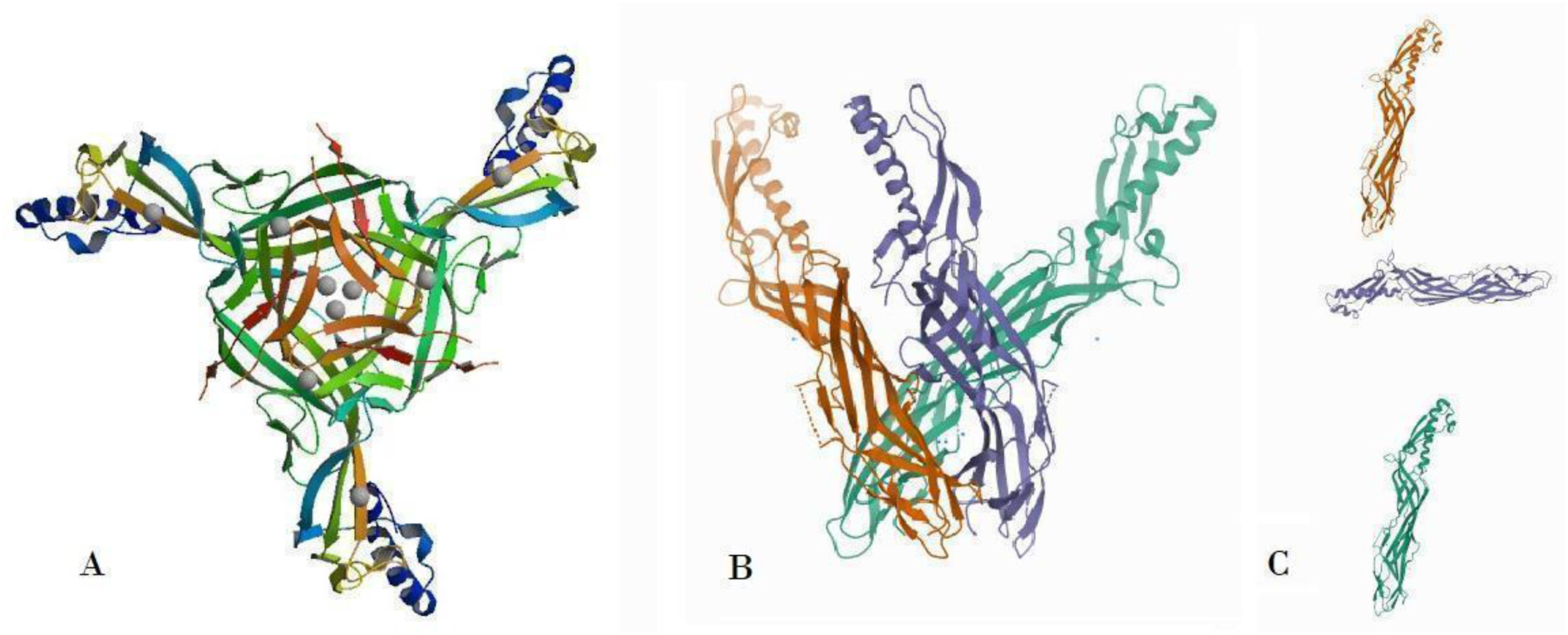
shows the 3D structure of *C. perfringens* Epsilon Toxin (ETX) was retrieved from the Protein Data Bank (PDB) with the PDB ID: 1UYJ (ETX-PDB). The structure provides insights into the spatial arrangement of ETX molecules. A and B represent the Biological Assemblies, depicting how the ETX chains interact and form larger structures. C corresponds to the Asymmetric Unit, illustrating the smallest repeating unit of the crystal structure.

For ATX, lacking a crystallography-derived 3D model, EasyModeller 4.0 generated a 3D model (ATX- 3D) through homology modelling (Figure 3). After energy minimization using the Swiss-Pdb Viewer 4.1.0 software, the Structure validation of ATX-3D was assessed via its Ramachandran plot from RAMPAGE (Figure 4). The plot shows 1887 (86.7%) residues in the favoured region, 188 (8.6%) in the allowed region, and 101 (4.6%) in the outlier region. With 95.3% of residues in the valid area, the ATX- 3D model can be considered ideal.

**Figure 3.**
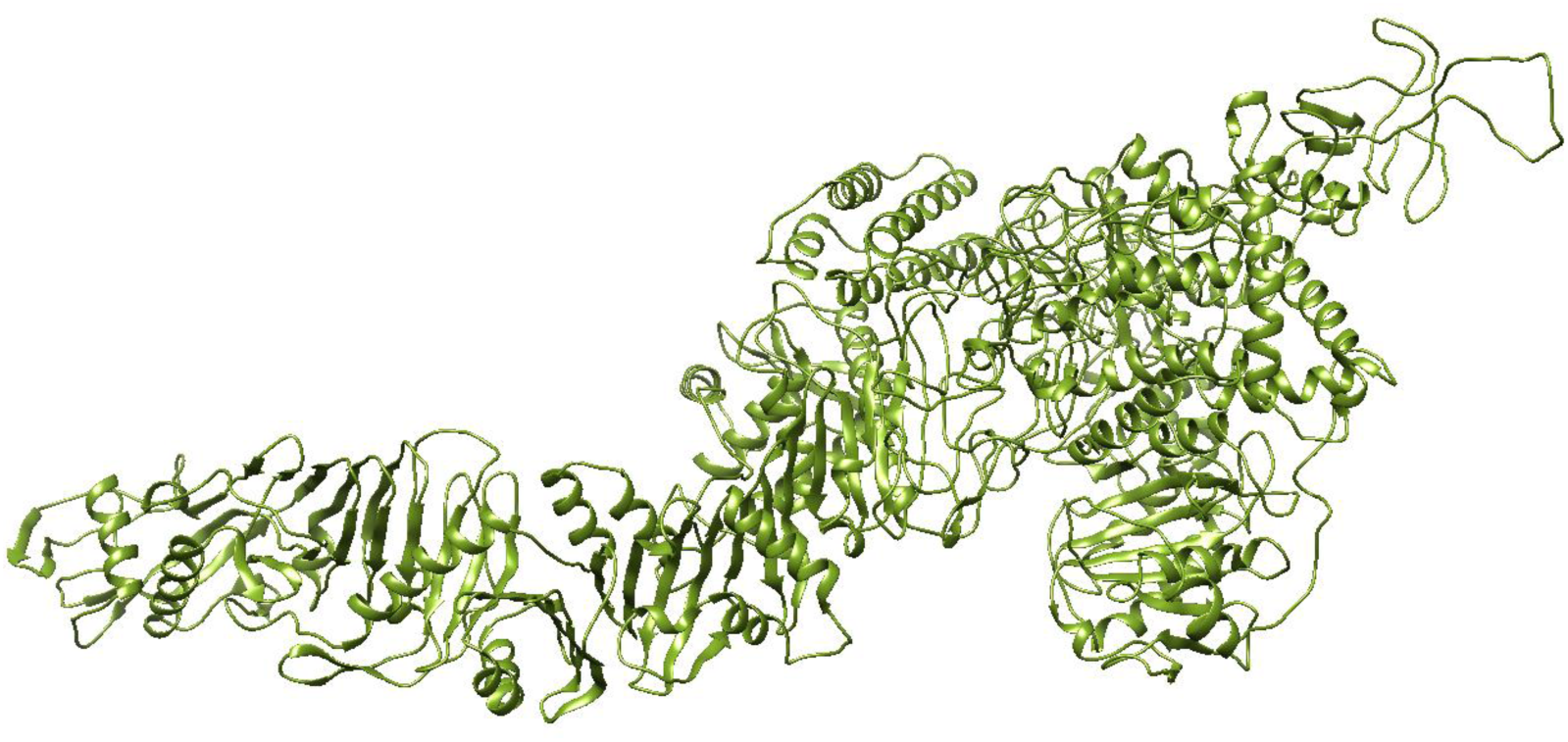
represents the *C. novyi* ATX 3D model predicted by EasyModeller 4.0 software using homology modelling (ATX-3D).

**Figure 4.**
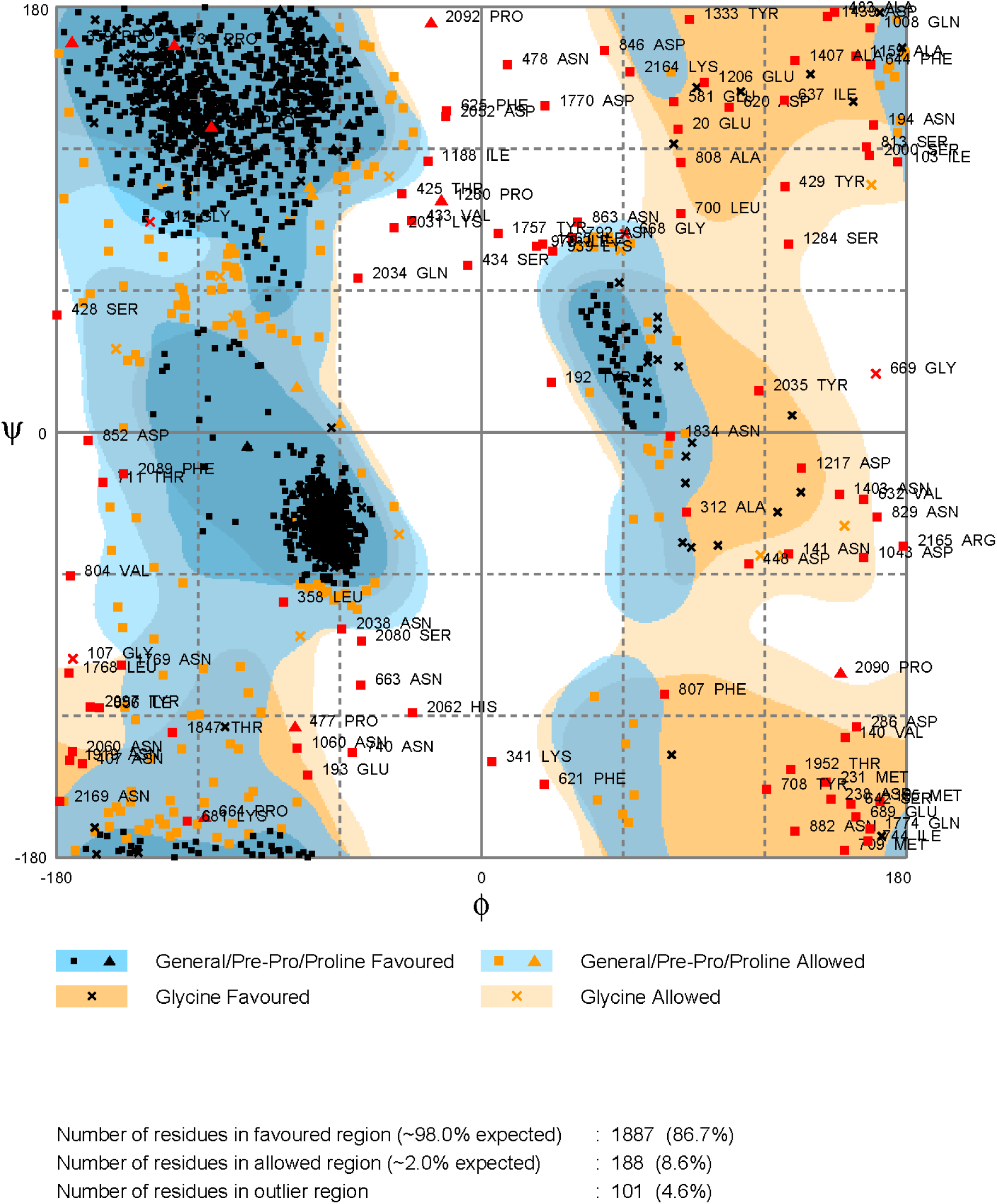
displays the Ramachandran plot for ATX-3D generated by RAMPAGE. This diagram illustrates the ϕ versus ψ backbone conformational angles in proteins. It showcases observed conformations or calculated energy contours for dipeptides. In recent years, these plots have become vital for structure validation, serving as a sensitive indicator of local issues since ϕ,ψ values are not optimized in the refinement process. The Ramachandran diagram, summarizing information on backbone conformation, has significantly contributed to our comprehension of protein structure, energetics, and folding. The plotindicates that 1887 residues (86.7%) fall within the favoured region, 188 residues (8.6%) in the allowed region, and 101 residues (4.6%) in the outlier region. Considering that 95.3% of residues lie in the valid area, the ATX-3D model can be deemed ideal (Lovell et al., 2003).

### 3.5. Conformational B-cell epitope identification

Conformational B-cell epitopes for ETX-PDB (excluding 1-32 residues as the signal peptide sequence) and ATX-3D (all 2178 residues) were predicted using three databases: CBTope (-0.3 threshold), DiscoTope-2.0-CBS (-0.37 and 0.5 thresholds), and Ellipro-IEDB (0.5 threshold) as shown in Table S4 and Table S5 (Supplementary material Appendix 1). Ellipro outputs were visually represented as 3D structures in Figures 5, 6, 7 and 8.

**Figure 5.**
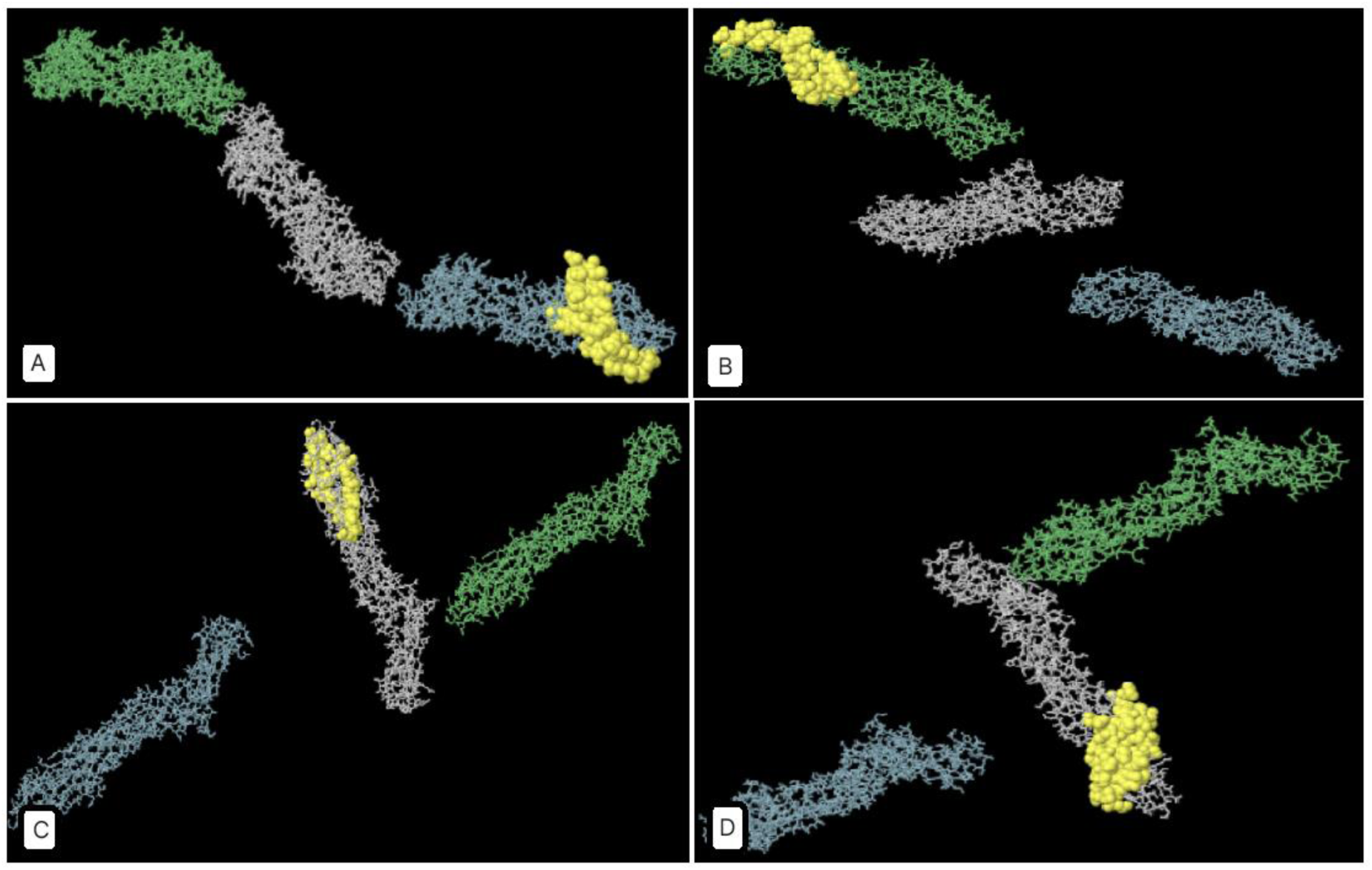
depicts a 3D representation of the highest-scoring Linear B-cell epitopes of *C. perfringens* ETX, as predicted by IEDB-Ellipro (highlighted in yellow). ElliPro, the prediction tool, utilizes ellipsoids to approximate the 3D structure of the protein antigen. Each epitope is associated with a Protrusion Index (PI) score, signifying the proportion of protein residues within a specific ellipsoid. Higher PI values imply a larger inclusion of residues in the epitope, indicating enhanced solvent accessibility and potential antigenicity. A: Chain A’s highest-scoring Linear B-cell epitope spans residues 246-295 (PI score: 0.786). B: Chain B’s highest-scoring Linear B-cell epitope encompasses residues 195-239 (PI score: 0.719). C and D: Chain C boasts two top-scoring Linear B-cell epitopes (C: Residues 159-184, PI score: 0.882 - D: Residues 250-295, PI score: 0.875). Additional details can be found in Supplementary material Appendix 1, Table S2.

**Figure 6.**
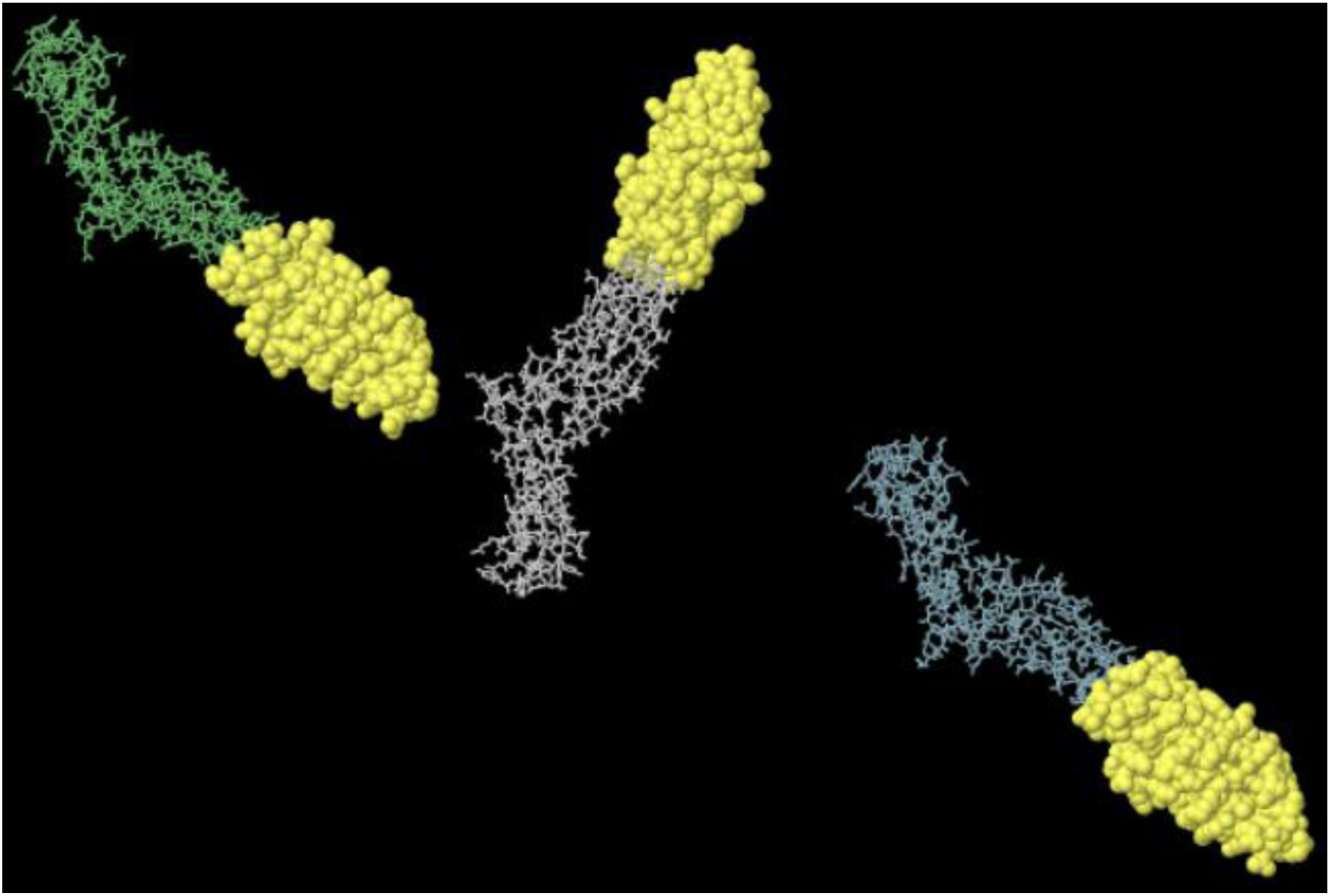
shows the 3D representation of the highest scored (PI score: 0.856) Discontinues B- cell epitope of *C. perfringens* ETX by IEDB - Ellipro (Details in Supplementary material Appendix 1, Table S4)

**Figure 7.**
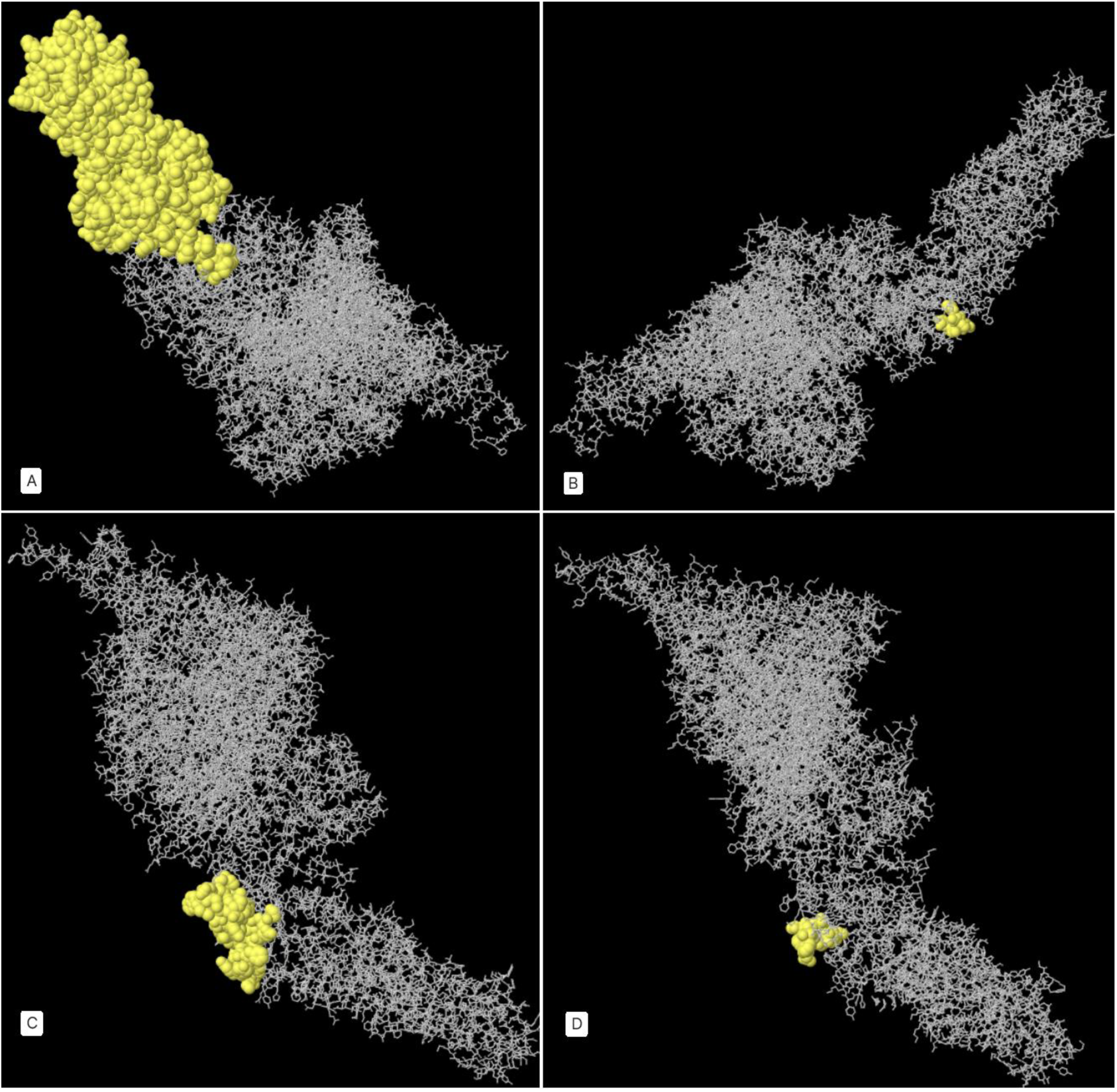
represents the 3D illustration of the high scored Linear B-cell epitopes of *C. novyi* ATX by IEDB - Ellipro. A: The top-scoring Linear B-cell epitope spans residues 1375-1697 (PI score: 0.839). B: A high-scoring Linear B-cell epitope encompasses residues 1846-1852 (PI score: 0.717). C: Another high-scoring Linear B-cell epitope is found in residues 1862-1907 (PI score: 0.695). D: A high-scoring Linear B-cell epitope is present in residues 1829-1839 (PI score: 0.688). (More details in Supplementary material Appendix 1, Table S4)

**Figure 8.**
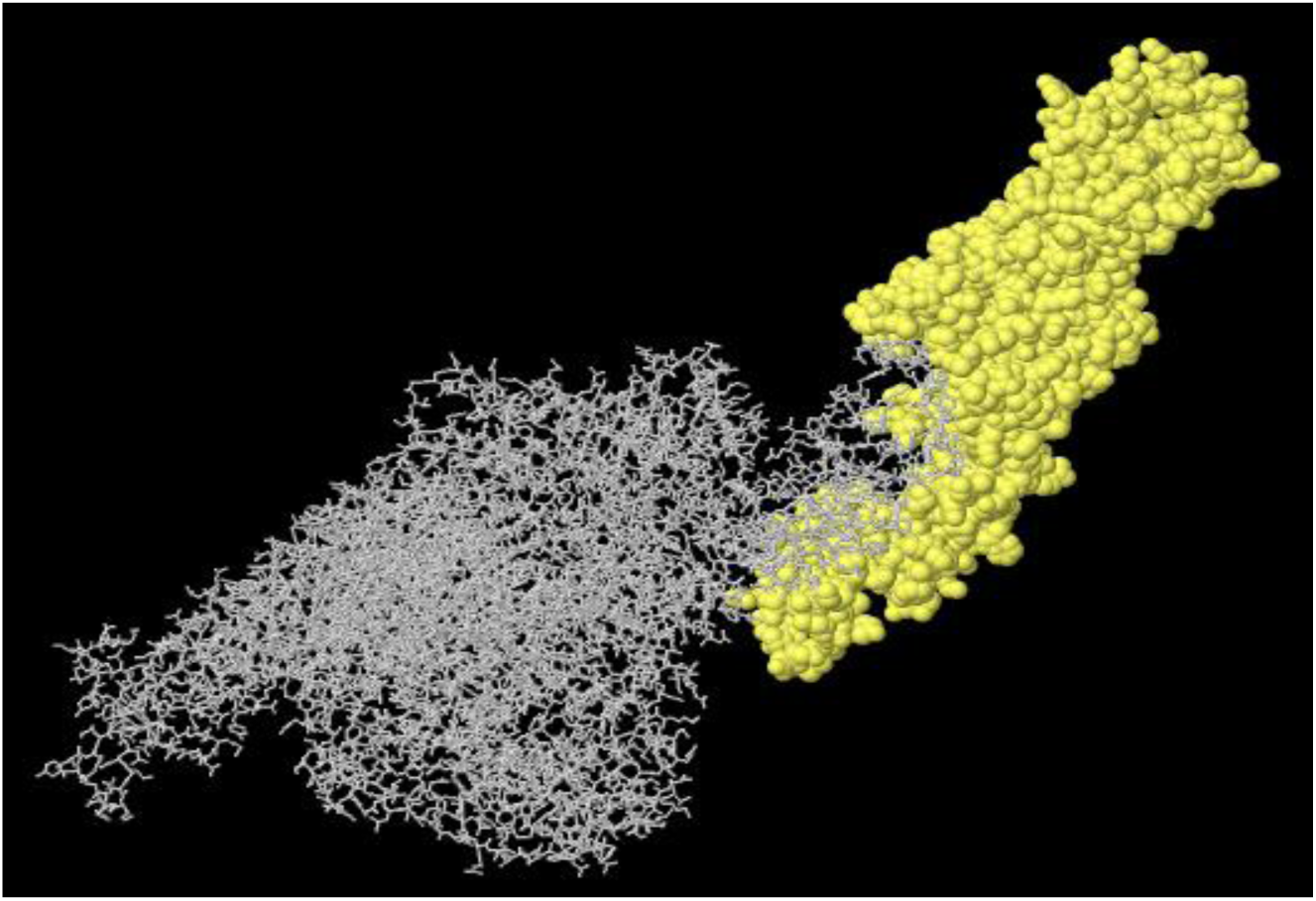
depicts the highest scored (472aa, PI score: 0.779) Discontinues B-cell epitope of *C. novyi* ATX by IEDB - Ellipro in yellow (Details in Supplementary material Appendix 1, Table S5).

Based on continuous and discontinuous B-cell epitope predictions, the most immunogenic epitopes, considering antigenicity and hydrophobicity plots along with the second structure of the toxins, were identified between residues 200-300 aa and 1800-2178 aa for ETX and ATX, respectively. The highly epitopic fragments ETXF (199-302 aa) and ATXF (1822-1992 aa) were chosen as the most antigenic regions of each toxin. The 3D molecular structures of ETXF and ATXF were visualized using ribbon and surface representations by UCSF Chimera 1.8 (Figure 9).

**Figure 9.**
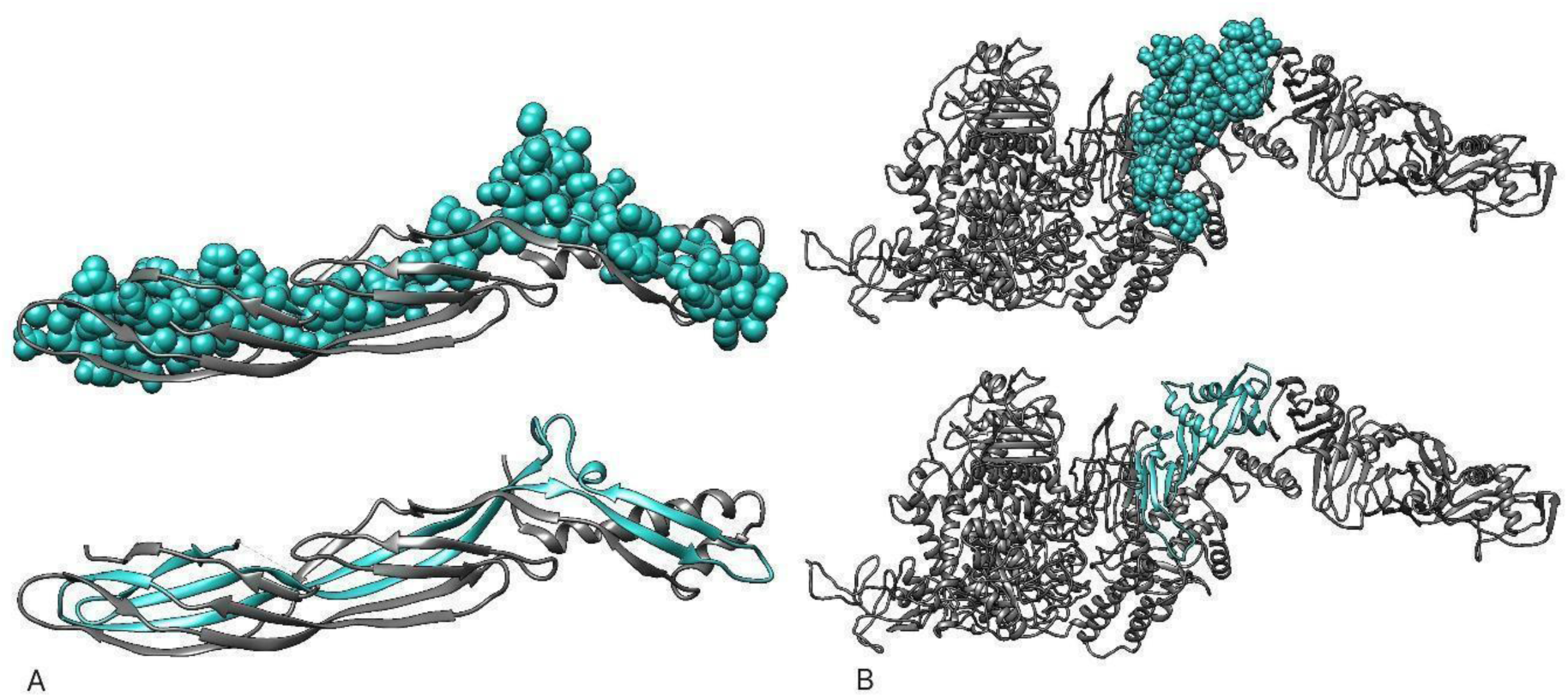
presents a 3D representation of the selected epitopic fragments of A: *C. perfringens* ETX (ETXF) and B: *C. novyi* ATX (ATXF) using UCSF Chimera 1.8. The ribbon and surface visualization provide a comprehensive depiction of the spatial arrangement and structural characteristics of the chosen epitopes. The ribbon representation highlights the backbone structure of the epitopes, offering insights into their overall conformation, while the surface visualization provides a detailed view of the molecular surface features. UCSF Chimera 1.8, a versatile molecular visualization tool, facilitates the exploration and analysis of these epitopic fragments in three dimensions, aiding in a more thorough understanding of their structural attributes.

### 3.6. Design of the recombinant fusion protein

To construct a recombinant fusion protein (rFP) incorporating ETXF, ATXF, E1, E2, A1, A2, and the PADRE peptide as an adjuvant, we assessed five distinct models (A-E). MHC-II binding epitopes (A1, E1, A2, E2) were combined to create a T-helper Epitopic Fragment (TEF), with the linker GPGPG used between each T-helper epitope. The epitopes were arranged in descending order of their scores to enhance the immune system response. To ensure proper folding of B-cell epitopic fragments within the native protein, we employed the GGSSGG linker. This linker not only facilitates correct folding but also serves as a target for proteases in the blood upon introduction of the vaccine candidate. As a result, targeted cleavage occurs rather than random cleavage. Models A-E were scrutinized as potential FP candidates (see Figure 10, and Supplementary material Appendix 1, Table S6).

**Figure 10.**
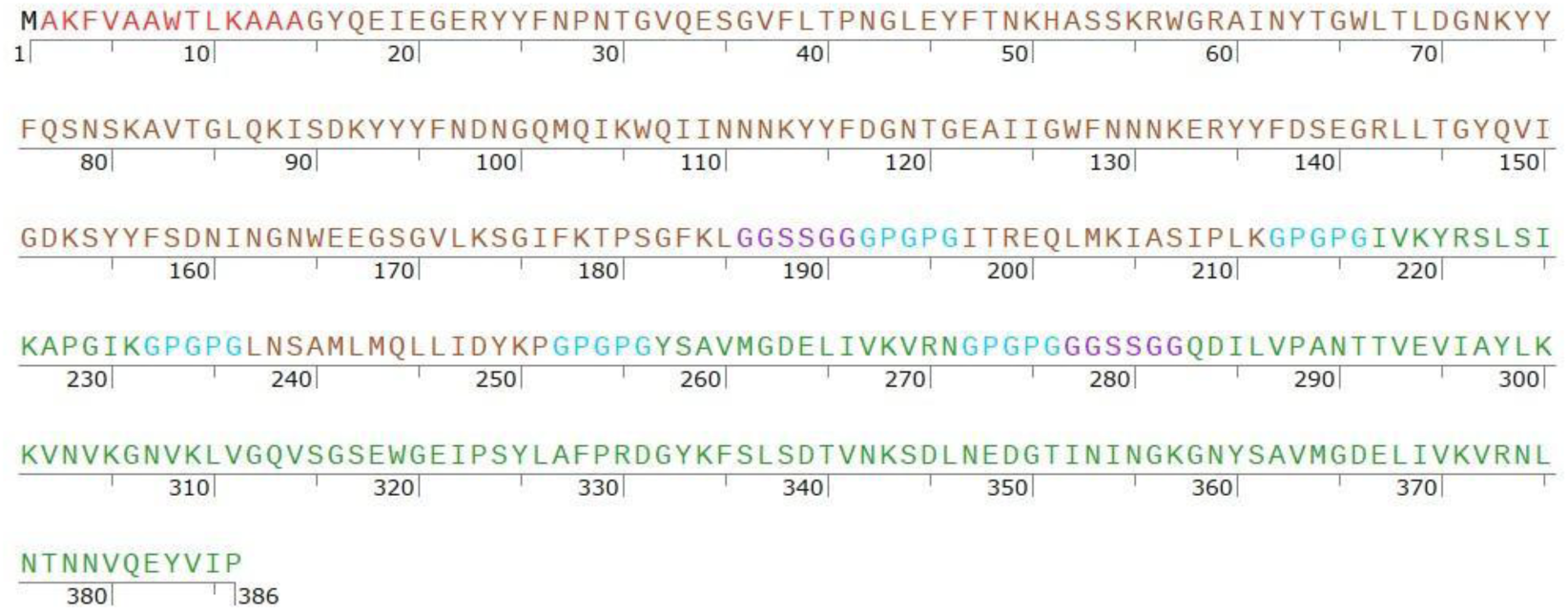
illustrates the amino acid sequence of the selected Recombinant Fusion Protein D (rFPD) derived from epitopic fragments and MHC-II binding epitopes of *C. perfringens* ETX and *C. novyi* ATX. The sequence consists of 386 amino acid residues and starts with methionine attached to the N-terminal end, serving as the initial amino acid. The adjuvant, PADRE sequence, is highlighted in red for clarity. TEF, representing T-helper Epitopic Fragment, encompasses regions A1, E1, A2, and E2, crucial for eliciting immune response. ATXF, denoting selected fragments from *C. novyi* ATX, is depicted in brown, along with A1 and A2, the highest-scoring Helper T-cell epitopes for ATX. Similarly, ETFX, selected fragments from *C. perfringens* ETX, and E1 and E2, the top-scoring Helper T-cell epitopes for ETX, are depicted in green. GGSSGG serves as the linker for B-cell epitopes, shown in purple, while the linker for Helper T-cell epitopes contains the GPGPG sequence, represented in blue.

EXPASY-ProtParam results reveal that Recombinant Fusion Protein D (rFPD), with a molecular weight of 42 kDa and an isoelectric pH of 9.09, exhibits a suitable half-life in vitro (30 hours) and in vivo (over 10 hours in E. coli). With an instability index of 20.71, rFPD is deemed a stable protein (instability index below 40). The GRAVY (Grand Average of Hydropathicity) index of rFPD is -0.443, indicating its immunogenic nature. These physicochemical properties for all the constructions were similar. However, in rFPD, ATX and ETX are far from each other to not influence each other’s structures. Therefore, rFPD was chosen as the most promising vaccine candidate against ETX and ATX (see Figure 1-F). Table S8 provides details of the protein sequence of rFPD (Supplementary material Appendix 1). AlgPred and AllergenFP v.1.0 outputs confirm that rFPD is non-allergenic.

### 3.7. Reverse translation of the Recombinant Fusion Protein D (rFPD)

The reverse translation process of the Recombinant Fusion Protein D (rFPD) involved several steps to ensure accurate reconstruction of the nucleotide sequence. First, the nucleotide sequences of the ETXF, ATXF, and TEF fragments were derived from the main toxins. Reverse translation of the sequences of linkers and the PADRE peptide was performed using BackTranseq-EMBOSS. For codon usage adaptation, CG content optimization, and restriction enzyme site analysis, the JCat server was employed. A start codon (ATG) was designated, and a stop codon (TAG) was appended to construct the *rfpd* gene, resulting in a total nucleotide sequence length of 1161 bp (Supplementary material Appendix 1, Table S9). Subsequently, the *rfpd* sequence underwent translation into a protein using CLC Main workbench 5.5. EXPASY-MultAlin verified that the reverse translation accurately reconstructed the rFPD protein from the *rfpd* gene.

The information obtained from CLC Main Workbench 5.5 regarding the presence of the several restriction enzyme recognition sequences within the *rfpd* gene is valuable for molecular cloning purposes. The identification of this sequence alerts us to avoid using these restriction enzymes for cloning the *rfpd* gene, as it could lead to unintended cleavage and disruption of the gene sequence during the cloning process. This insight helps in selecting appropriate restriction enzymes for cloning that will not interfere with the integrity and functionality of the *rfpd* gene construct (Figure 11).

**Figure 11.**
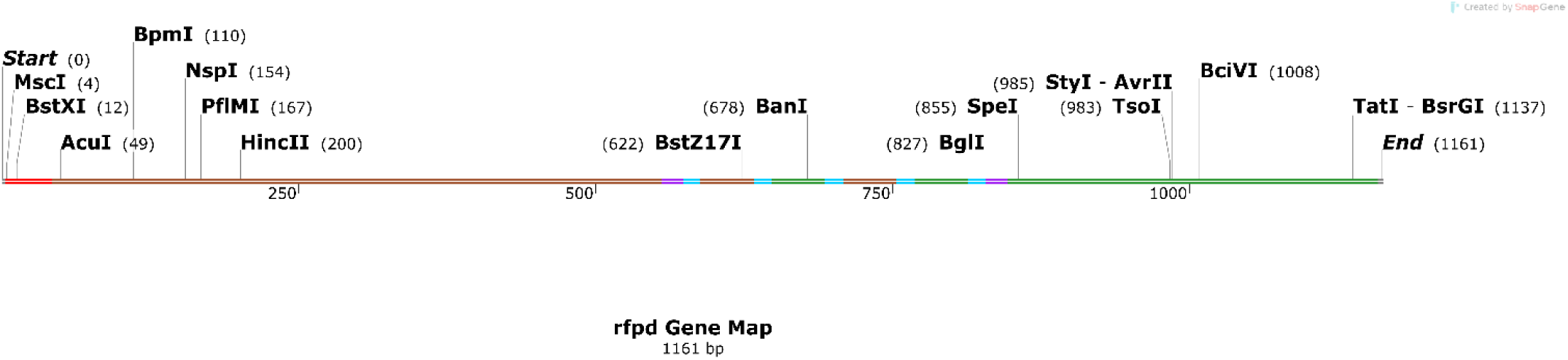
presents the reverse translation of the Recombinant Fusion Protein D (rFPD) into the corresponding nucleotide sequence, referred to as the *rfpd* gene. The nucleotide sequence of the *rfpd* gene comprises 1161 base pairs (bp). The initiation codon “ATG” signifies the start codon located at the N-terminal end, initiating the translation process, while the termination codon “TAG” denotes the stop codon positioned at the C-terminal end, signifying the end of protein synthesis. Highlighted in red, the adjuvant PADRE sequence is prominently displayed for clarity, indicating its pivotal role in enhancing immunogenicity. Additionally, the table delineates specific regions within the *rfpd* gene, such as the T-helper Epitopic Fragment (TEF), comprising regions A1, E1, A2, and E2, crucial for eliciting an immune response. Moreover, the table categorizes selected fragments from *C. novyi* ATX, as ATXF, depicted in brown, along with the highest-scoring Helper T-cell epitopes (A1 and A2) for ATX. Similarly, ETFX, selected fragments from *C. perfringens* ETX, denoted as ETFX, are represented in green, along with the top-scoring Helper T-cell epitopes (E1 and E2) for ETX. The nucleotide sequence of the linkers for B-cell epitopes depicted in purple, facilitating the structural integrity and functionality of the fusion protein. The sequence for the linkers of Helper T-cell epitopes is represented in blue, ensuring proper spatial arrangement and interaction with immune cells. There are several restriction enzyme sites within the *rfpd* gene that assist in the selection of appropriate restriction enzymes for future cloning purposes.

## 4. Discussion

*Clostridium perfringens* type D causes fatal enteric diseases, while *Clostridium novyi* type B is responsible for lethal gangrene in domestic ruminants. The key virulence factors in these diseases are *C. perfringens* type D Epsilon toxin (ETX) and *C. novyi* type B Alpha toxin. (ATX) (Hatheway, 1990) Given the high fatality rate associated with both bacteria, there is insufficient time for antibiotic therapy. Consequently, the most viable alternative is to vaccinate animals against these infections.

Conventional vaccines, especially toxoids, are hindered by challenges such as undesirable side effects, non-specific immune responses, and the cumbersome, costly manufacturing process linked to anaerobic bacteria. To address these issues, contemporary approaches in immune informatics have been employed to explore novel immunogenic regions and epitopes within proteins of interest. This innovative strategy aims to elicit specific and targeted immune responses, ultimately contributing to cost-effective production methods for diverse bacterial organisms.

In this study, various immunoinformatic tools were employed to design and express a novel Recombinant Fusion Protein (rFPD) serving as a vaccine candidate against both *C. perfringens* type D and *C. novyi* type B simultaneously. rFPD comprises the optimal epitopic fragments of ETX (ETXF: 199- 302 aa) and ATX (ATXF: 1822-1992 aa), along with the two most immunogenic T-helper epitopes from both toxins (E1, E2, A1, A2). The selection of these epitopes was based on results from T-helper and B-cell epitope databases, consideration of their secondary structures, and analysis of hydrophobicity and antigenicity plots of the toxins. Additionally, the inclusion of the PADRE peptide (AKFVAAWTLKAAA) as an adjuvant aimed to enhance the immunogenicity of rFPD. Proper linkers were strategically utilized to connect the fragments, preventing the formation of new epitopes, interference between selected epitopes, and issues related to immunodominance and fragments.

The choice of antigenic determinants in vaccine design is crucial for inducing effective immune responses. Previous studies have indicated that coil structures on the molecule’s surface and β-sheets are more likely to serve as antigenic determinants, eliciting higher immune responses compared to α- helix structures (Ghaffari-Nazari et al., 2015; Modrow et al., 1987) In line with these findings, the selection of epitopic fragments ETXF and ATXF from loop-like and β-sheet regions, as revealed by immunoinformatic databases, aligns with the goal of maximizing the vaccine’s antigenicity. Moreover, the decision to choose fragments based on overlapping continuous and conformational B-cell epitope regions adds an extra layer of promise to these fragments as highly antigenic components.

The confirmation of our bioinformatics analysis by the inclusion of highly antigenic continuous B-cell epitopes aligns with the findings of a recent study by Alves et al. (Alves et al., 2017). This study identifies four highly antigenic continuous B-cell epitopes out of 15 potential antigenic determinants for developing subunit vaccines against *C. perfringens* type D ETX. Notably, ETXF, which we selected as the antigenic fragment for ETX in our design, encompasses three of these confirmed epitopic determinants. This region’s involvement in toxin attachment to the membrane and its role in the interaction between the three ETX chains to form the cell pore emphasizes its significance. The presence of antibodies against this region may potentially prevent the activation of ETX by inhibiting the cleavage of the protoxin by enzymes.

The alignment of our selected fragment, ATXF (1822-1992 aa), with the highly antigenic C-terminal segment of ATX, specifically pNTF12 (1800-1958 aa), further supports the rationale behind our immunoinformatic design. According to research by Busch et al. and Gord-Noshahri et al. (Busch et al., 2000; Gord-Noshahri et al., 2016), pNTF12 is identified as a hydrophilic surface protein responsible for receptor binding activity. It exhibits high mobility and lacks catalytic activity. Notably, our chosen ATXF fragment substantially overlaps with the critical pNTF12 region.

Gord-Noshahri et al. (Gord-Noshahri et al., 2016) demonstrated active recognition of pNTF12 by anti-ATX antibodies in immunological assays. Anti-NTF12 antibodies produced in mice not only recognized ATX but also exhibited higher reactivity to native ATX compared to anti-ATX antibodies. This observation suggests that the 3D structure and epitopes of pNTF12, and consequently ATXF, are likely to be retained in their native form.

## 5. Conclusions

In conclusion, our in-silico design of a vaccine candidate, the Recombinant Fusion Protein D (rFPD), holds the potential for providing simultaneous immunity against both *Clostridium perfringens* type D and *Clostridium novyi* type B. rFPD, a fusion peptide or polyepitopic construct, has been meticulously designed using immunoinformatic tools, incorporating the best epitopic fragments of Epsilon toxin (ETX) and Alpha toxin (ATX), along with two highly immunogenic T-helper epitopes from each toxin (E1, E2, A1, A2). The inclusion of the PADRE peptide as an adjuvant further enhances the immunogenicity of rFPD.

The next crucial steps involve the synthesis and evaluation of rFPD through animal experiments, where the effectiveness and protective efficacy of the immunization will be assessed in comparison with existing vaccines. These experiments will provide valuable insights into the potential of rFPD as a promising vaccine candidate against infections caused by *C. perfringens* type D and *C. novyi* type B.

