## Supplementary file for "In Silico Vaccine Design: Targeting Highly Epitopic Regions of *Clostridium perfringens* Type D Epsilon Toxin and *Clostridium novyi* Type B Alpha Toxin for Optimal Immunogenicity"

| Toxin | Start | End | Peptide | Percentile rank in IEDB |
|---|---|---|---|---|
| ETX | 314 | 328 | E1: IVKYRSLSIKAPGIK | 0.50 |
| | 276 | 290 | E2: YSAVMGDELIVKVRN | 9.46 |
| ATX | 3 | 17 | A1: ITREQLMKIASIPLK | 0.08 |
| | 965 | 979 | A2: LNSAMLMQLLIDYKP | 0.93 |

**Table S1** indicates the two highest scored MHC-II binding epitopes(H2-IAd) for *C. perfringens* type D Epsilon Toxin, ETX, (E1, E2) and (A1, A2) for *C. novyi* type B Alpha Toxin, ATX. The predictions were made using the Immune Epitope Database (IEDB) server with the Consensus (smm/nn) method. A lower percentile rank in IEDB indicates higher immunogenicity of the epitope.

| Server name | Sequence (aa) |
|---|---|
| LBTope | 39-46, 102-112, **105**, **108**, 128-134,184-187, 193-194, 251-255, 262-266, 273, 280-303, **290**, **293, 295** |
| SVMTriP | **99-118**, **143-162**, 205-224, 57-76 |
| BepiPred-2.0 (CBS) | 33-61, 67-94, 101-111, 125-138, 179-183, 193-201, 239-265, 291-299, 301-324 |
| ABCpred (0.6: epitope threshold) (16 aa length) | 1: **84-99, 104, 154, 170**<br>69, 278, 142, 208, 265, 61, 296, 202, 52, 235, 242, 12, 179, 189, 39, 115, 134, 93, 161, 127, 310, 251, 195, 121 |
| Bepipred (0.6) | 38, 40-50, 61-62, 87-94, 101-110, 125-132, 142-148, 150-152, 158, 172, 181-197, 231-237, 257-265, 267-271, 273-275, 304-312 |
| Kloskar (1.0) | 81-88, 112-123, 130-139, 150-156, 171-180, 193-206, 208-232, 238-245, 250-256, 281-289, 297-304 |

| | |
|---|---|
| Emini surface accessibility (1.0) | 42-48, 58-76, 124-133, 183-192, 304-310 |
| Ellipro (Linear Epitope) | 257-265, 221-237, 32-66, 197-218, 83-108, 271-295, 159-178 |

**Table S2** presents the linear B-cell epitopes of *C. perfringens* ETX. Amino acid (aa) sequences display highly immunogenic B-cell epitopes with a probability range of 61-80%. The bolded sequences represent B-cell epitopes with a higher probability, ranging from 81-100%. Highlighted sequences indicate the main epitopic region for ETX, located between residues 150-300.

| Server name | Sequence (aa) |
|---|---|
| LBTope | 6-11, 17-24, 59-62, 90-97, 157-163, 181-188, 278-289, **283-284** 327-339, **333**, **336**, **338**, 359-369, 613-623, 729-741, 860-865, 915-939, **923**, **924**, 953-965, **957-961**, 1058-1062, 1182-1189, 1215-1225, 1270-1288, **1282**, 1343-1364, **1345**, 1436-1452, **1356**, 1466-1470, 1500-1512, **1509-1510**, 1500-1524, 1564-1574, 1634-1646, **1636-1637**, 1780-1785, 1789-1800, **1791-1794**, 1854-1867, **1858-1862**, 1881-1884, 1898-1899, 1923-1929, 2003-2005, 2073-2075 |
| SVMTriP | 23-32, 1419-1428, 383-392, 292-301, 1040-1051 |
| BepiPred-2.0 (CBS) (0.6) | 576-578, 632-643, 736-737, 848-850, 1184-1190, 1214-1215, 1248, 1648-1654, 1970-1974 |
| ABCpred (0.6: epitope threshold) (16 aa length) | 0.9> 1562, 1432, 914, 296, 1685, 1618, 1257, 1202, 922, 893, 278, 2109, 1999, 1823, 1710, 1674<br>0.85> 705, 2100, 1856, 1595, 455, 419, 2031, 1912, 1396, 1001, 947, 537, 2073, 1872, 1652, 152, 1289, 84, 1438, 1306 |
| Bepipred (-0.185) (Size) | 1820-1869 (50), 2048-2096(49), 1874-1898(25), 1955-1979(25), 2007-2031(25), 153-175(23), 1232-1254(23), 593-613(21), 1983-2003(21), 322-341(20), 1264-1283(20), 729-747(19), 908-926(19), 1648-1666(19), 659-674(16), 1340-1355(16), 56-70(15), 438-452(15), 461-475(15) |
| Kloskar (1.0) 0.9> | 118-126, 193-200, 221-233, 237-243, 300-309, 555-561, 752-760, 816-822, 881-888, 1247-1253, 1266-1272, 1314-1322, 1349-1355, 1622-1629, 1724-1730, 1903-1910, 1923-1930, 1933-1942, 1965-1977 |
| Emini surface accessibility (1.0) 0.2> | 11-16, 25-30, 100-109, 201-210, 231-237, 269-275, 280-288, 313-319, 353-361, 368-373, 385-392, 411-416, 433-438, 456-461, 601-606, 613-618, 702-710, 720-725, 860-865, 871-879, 887-896, 938-946, 970-975, 999-1007, 1012-1021, 1036-1052, 1073-1088, 1084-1094, 1096-1113, 1115-1124, 1180-1189, 1223-1230, 1240-1245, 1297-1302, 1307-1315, 1334-1340, 1372-1378, 1416-1425, 1432-1438, 1457-1462, 1466-1472, 1578-1591, 1589-1603, 1619-1624, 1640-1648, 1673-1682, 1687-1692, 1709-1718, 1739-1744, 1746-1751, 1814-1819, 1928-1936 |
| Ellipro (Linear Epitope) | 889-955, 1375-1679, 186-282, 505-565, 971-1000, 840-882, 1846-1852, 1776-1813, 1862-1907 |

**Table S3** shows the Linear B-cell epitopes of *C.novyi* ATX. Amino acid (aa) sequences display highly immunogenic B-cell epitopes with a probability range of 61-80%. The bolded sequences represent B-cell epitopes with a higher probability, ranging from 81-100%. Highlighted epitopes show that the main epitopic region for ATX is between 1200-2178 residues.

| Server name | | Sequence (aa) + 32 |
|---|---|---|
| CBTope | | 54-62, 67-69, 74, 82-93, **89-108**, 112-115, 118-123, 137,139, 169, 170-177, 183, 195-201, 212, 215-216, 242-251, **253-278**, 280-283, 288-290, 300, 302, 318-323 |
| DiscoTope -2.0- CBS (-2) | A | 16-17, 20, 23, 25-61, 63, 74, 93-96, 123, 126, 173-174, 199-220, 222-223, 226-235 |
| | B | 22-23, 25-61, 63, 74, 93-96, 123, 173-174, 197, 199-220, 222-223, 226-235, 237 |
| | C | 23, 25-61, 63, 74, 93-96, 123, 126, 173-174, 199-223, 225-235, 237 |
| Ellipro (Discontinus) | | > 101aa, score: 0.856<br>C:Y79, C:L80, C:E81, C:D82, C:V83, C:Y84, C:V85, C:G86, C:K87, C:A88, C:L89, C:L90, C:T91, C:N92, C:D93, C:T94, C:Q95, C:Q96, C:E97, C:Q98, C:K99, C:L100, C:K101, C:S102, C:Q103, C:S104, C:F105, C:T106, C:C107, C:K108, C:N109, C:N136, C:E137, C:T138, C:G139, C:V140, C:E159, C:I160, C:T161, C:H162, C:N163, C:V164, C:P165, C:S166, C:Q167, C:D168, C:I169, C:L170, C:V171, C:P172, C:A173, C:N174, C:T175, C:T176, C:V177, C:E178, C:V179, C:I180, C:A181, C:Y182, C:L183, C:K184, C:D250, C:E251, C:L252, C:I253, C:V254, C:K255, C:V256, C:R257, C:N258, C:L259, C:N260, C:T261, C:N262, C:N263, C:V264, C:Q265, C:E266, C:Y267, C:V268, C:I269, C:P270, C:V271, C:D272, C:S280, C:N281, C:I282, C:V283, C:K284, C:Y285, C:R286, C:S287, C:L288, C:S289, C:I290, C:K291, C:A292, C:P293, C:G294, C:I295 |
| Ellipro (3D-Linear) | 1 | > A chain, 246aa-295aa (43aa), score: 0.786<br>AVMGDELIVKVRNLNTNNVQEYVIPVDSNIVKYRSLSIKAPGI |
| | 2 | > B chain, 195aa-239aa (45aa), score: 0.719<br>VGQVSGSEWGEIPSYLAFPRDGYKFSLSDTVNKSDLNEDGTININ |
| | 3 | > C chain, 159aa-184aa (26aa), score: 0.882<br>EITHNVPSQDILVPANTTVEVIAYLK |
| | 4 | > C chain, 259aa-295aa (39aa), score: 0.875<br>DELIVKVRNLNTNNVQEYVIPVDSNIVKYRSLSIKAPGI |

**Table S4.** Conformational B-cell epitopes of *C. perfringens* ETX. Highlighted epitopes show that the main epitopic region for ETX is between 150-300 residues. Signal peptide is eliminated.

| Server name | Sequence (aa) |
|---|---|
| CBTope | 27, 55, 58-75, 77, 99-100, 102, 104-109, 115-122, 126, 128-130, 155-179, 193-199, 201, 284-285, 287, 289, 291-299, 302-306, 311-314, 329-333, 336, 339-346, 387-389, 410-413, 429, 445-447, 449-470, 476-482, 486-487, 489-495, 517, 519-521, 568-569, 584-586, 590, 593-594, 598-599, 608, 615, 631-632, 639, 655-656, 662, 665, 672-674, 676-677, 682-683, 685-686, 705-710, 724-725, 733-740, 759-765, 772, 775, 790-791, 811-814, 816, 818-828, 839, 849-853, 894-895, 905-907, 921-924, 927-928, 931, 985, 989, 996-997, 1044-1050, 1083- |

| | | |
|---|---|---|
| | 1089, 1144-1145, 1173, 1224, 1227, 1240-1242, 1246, 1256-1257, 1267-1268, 1270, 1272, 1277-1281, 1301-13022, 1304-1309, 1335-1339, 1344-1351, 1360, 1363-1365, 1368-1369, 1372, 1395-1398, 1402-1407, 1430-1431, 1436-1438, 1470-1472, 1474-1488, 1524-1537, 1540-1544, 1572-1573, 1602-1603, 1608-1614, 1616,1620, 1630-1633, 1636-1646, 1648-1651, 1653-1654, 1659-1668, 1671, 1735-1739, 1747, 1810, 1842-1845, 1847-1851, 1853-1858, 1861-1863, 1865-1868, 1873-1875, 1879-1905, 1907-1912, 1914-1916, 1920-1941, 1943, 1951-1958, 1961, 1965-1968, 1970-1984, 1986-1991, 2004-2005, 2007-2017, 2019-2029, 2053-2061, 2071-2080, 2082, 2101-2120, 2129-2140, 2169-2178 | |
| DiscoTope - 2.0- CBS (-2) | 14-19, 21, 63-65, 93, 125, 154, 157-58, 160-162, 164, 197-231, 236, 242-258, 260, 275, 277, 421-422, 494, 519, 534-535, 538, 541-542, 545-546, 548-552, 556, 558, 718, 860, 864, 866, 868-869, 871-873, 876-877, 924-926, 933, 935, 937, 1059, 1071, 1125-1126, 1161-1162, 1164, 1204, 1215, 1218, 1241, 1247-1261, 1264, 1268, 1271-1272, 1275, 1278-1280, 1321, 1323, 1325, 1327-1336, 1338, 1340-1344, 1346-1348, 1351, 1355-1358, 1360-1362, 1364-1365, 1367-1368, 1370, 1396-11398, 1400, 1442-1449, 1479, 1481-1482, 1488, 1493, 1501, 1521-1522, 1574, 1601, 1606, 1617, 1624, 1633-1635, 1637, 1639, 1641-1645, 1648, 1663-1665, 1678-1687, 1689-1690, 1692-1703, 1723-1728, 1790-1792, 1806, 1808-1809, 1811-1814, 1821-1822, 1827-1872, 1877-1882, 1885, 1889, 1892-1901, 1903, 1905-1925, 1927-1930, 1932-1948, 1953-1982, 1984-1985, 1988-1989, 1991-1995, 1997-1998, 2002-2009, 2011-2012, 2014-2015, 2045-2047, 2055, 2059, 2069-2074, 2105-2114, 2119-2126, 2148 | |
| Ellipro (Discontinus) | 1 | > 472aa, score: 0.779 _:D1375, _:G1376, _:F1377, _:I1378, _:N1379, _:I1381, _:F1382, _:S1383, _:T1384, _:L1385, _:K1386, _:V1387, _:S1388, _:N1389, _:D1390, _:G1391, _:F1392, _:K1393, _:I1394, _:G1395, _:K1396, _:Q1397, _:F1398, _:I1399, _:S1400, _:I1401, _:K1402, _:N1403, _:T1404, _:P1405, _:R1406, _:A1407, _:I1408, _:N1409, _:L1410, _:S1411, _:F1412, _:K1413, _:I1414, _:N1415, _:N1416, _:N1417, _:I1418, _:V1419, _:I1420, _:V1421, _:S1422, _:I1423, _:Y1424, _:L1425, _:N1426, _:H1427, _:E1428, _:K1429, _:S1430, _:N1431, _:S1432, _:I1433, _:T1434, _:I1435, _:I1436, _:S1437, _:S1438, _:D1439, _:L1440, _:N1441, _:D1442, _:I1443, _:K1444, _:N1445, _:N1446, _:F1447, _:D1448, _:N1449, _:L1450, _:L1451, _:D1452, _:N1453, _:I1454, _:N1455, _:Y1456, _:I1457, _:G1458, _:L1459, _:G1460, _:S1461, _:I1462, _:S1463, _:D1464, _:N1465, _:T1466, _:I1467, _:N1468, _:C1469, _:I1470, _:V1471, _:R1472, _:N1473, _:D1474, _:E1475, _:V1476, _:Y1477, _:M1478, _:E1479, _:G1480, _:K1481, _:I1482, _:F1483, _:L1484, _:N1485, _:E1486, _:K1487, _:K1488, _:L1489, _:V1490, _:F1491, _:I1492, _:Q1493, _:N1494, _:E1495, _:L1496, _:E1497, _:L1498, _:H1499, _:L1500, _:Y1501, _:D1502, _:S1503, _:V1504, _:N1505, _:K1506, _:D1507, _:S1508, _:Q1509, _:Y1510, _:L1511, _:I1512, _:N1513, _:N1514, _:P1515, _:I1516, _:N1517, _:N1518, _:V1519, _:V1520, _:K1521, _:Y1522, _:K1523, _:D1524, _:G1525, _:Y1526, _:I1527, _:V1528, _:E1529, _:G1530, _:T1531, _:F1532, _:L1533, _:I1534, _:N1535, |

| | | |
|---|---|---|
| | | _:S1536, _:T1537, _:E1538, _:N1539, _:K1540, _:Y1541, _:S1542, _:L1543, _:Y1544, _:I1545, _:E1546, _:N1547, _:N1548, _:K1549, _:I1550, _:M1551, _:L1552, _:K1553, _:G1554, _:L1555, _:Y1556, _:L1557, _:E1558, _:S1559, _:S1560, _:V1561, _:F1562, _:K1563, _:T1564, _:I1565, _:Q1566, _:D1567, _:K1568, _:I1569, _:Y1570, _:S1571, _:K1572, _:E1573, _:K1574, _:V1575, _:N1576, _:D1577, _:Y1578, _:I1579, _:L1580, _:S1581, _:L1582, _:I1583, _:K1584, _:K1585, _:F1586, _:F1587, _:T1588, _:V1589, _:N1590, _:I1591, _:Q1592, _:L1593, _:C1594, _:P1595, _:F1596, _:M1597, _:I1598, _:V1599, _:S1600, _:G1601, _:V1602, _:D1603, _:E1604, _:N1605, _:N1606, _:R1607, _:Y1608, _:L1609, _:E1610, _:Y1611, _:M1612, _:L1613, _:S1614, _:T1615, _:N1616, _:N1617, _:K1618, _:W1619, _:I1620, _:I1621, _:N1622, _:G1623, _:G1624, _:Y1625, _:W1626, _:E1627, _:N1628, _:D1629, _:F1630, _:N1631, _:N1632, _:Y1633, _:K1634, _:I1635, _:V1636, _:D1637, _:F1638, _:E1639, _:K1640, _:C1641, _:N1642, _:V1643, _:I1644, _:V1645, _:S1646, _:G1647, _:S1648, _:N1649, _:K1650, _:L1651, _:N1652, _:S1653, _:E1654, _:G1655, _:D1656, _:L1657, _:A1658, _:D1659, _:T1660, _:I1661, _:D1662, _:V1663, _:L1664, _:D1665, _:K1666, _:D1667, _:L1668, _:E1669, _:N1670, _:L1671, _:Y1672, _:I1673, _:D1674, _:S1675, _:V1676, _:I1677, _:I1678, _:I1679, _:P1680, _:V1682, _:Y1683, _:T1684, _:K1685, _:I1687, _:I1688, _:I1689, _:H1690, _:P1691, _:I1692, _:P1693, _:N1694, _:N1695, _:P1696, _:Q1697, _:I1698, _:N1699, _:I1700, _:I1701, _:Q1704, _:K1709, _:C1710, _:H1711, _:L1712, _:I1713, _:I1714, _:D1715, _:S1716, _:V1717, _:L1718, _:I1733, _:T1734, _:N1735, _:G1736, _:L1737, _:D1738, _:I1739, _:N1740, _:I1741, _:R1742, _:I1743, _:L1744, _:Q1745, _:G1746, _:L1747, _:S1748, _:F1749, _:G1750, _:F1751, _:K1752, _:Y1753, _:K1754, _:N1755, _:I1756, _:Y1763, _:D1764, _:E1765, _:L1766, _:S1767, _:L1768, _:N1769, _:D1770, _:F1771, _:L1772, _:L1773, _:Y1776, _:G1780, _:L1781, _:Y1782, _:Y1783, _:I1784, _:N1785, _:G1786, _:E1787, _:L1788, _:H1789, _:Y1790, _:N1792, _:I1793, _:P1794, _:G1795, _:D1796, _:T1797, _:F1798, _:E1799, _:Y1800, _:G1801, _:W1802, _:I1803, _:N1804, _:I1805, _:D1806, _:S1807, _:R1808, _:W1809, _:Y1810, _:F1811, _:F1812, _:D1813, _:G1822, _:E1829, _:R1830, _:Y1831, _:Y1832, _:F1833, _:N1834, _:P1835, _:N1836, _:T1837, _:G1838, _:V1839, _:L1846, _:T1847, _:P1848, _:N1849, _:G1850, _:L1851, _:E1852, _:S1860, _:K1862, _:R1863, _:G1865, _:R1866, _:A1867, _:N1869, _:Y1870, _:T1871, _:G1872, _:W1873, _:L1874, _:T1875, _:L1876, _:D1877, _:G1878, _:N1879, _:K1880, _:Y1881, _:Y1882, _:F1883, _:Q1884, _:S1885, _:N1886, _:S1887, _:K1888, _:A1889, _:V1890, _:T1891, _:G1892, _:L1893, _:Q1894, _:K1895, _:I1896, _:S1897, _:D1898, _:K1899, _:Y1900, _:Y1901, _:Y1902, _:F1903, _:N1904, _:D1905, _:N1906, _:G1907, _:G1928, _:E1929, _:A1930, _:G1982 |
| Ellipro (3D-Linear) | 1 | > 1375-1679 (305aa), score: 0.839<br>DGFINNIFSTLKVSNDGFKIGKQFISIKNTPRAINLSFKINNNIVIVSIYLNHE<br>KSNSITIISSDLNDIKNNFDNLLDNINYIGLGSISDNTINCIVRNDEVYMEG<br>KIFLNEKKLVFIQNELELHLYDSVNKDSQYLINNPINNVVKYKDGYIVEGTF<br>LINSTENKYSLYIENNKIMLKGLYLESSVFKTIQDKIYSKEKVNDYILSLIKKFF |

| | | |
|---|---|---|
| | | TVNIQLCPFMIVSGVDENNRYLEYMLSTNNKWIINGGYWENDFNNYKI VDFEKCNVIVSGSNKLNSEGDLADTIDVLDKDLENLYIDSVIII |
| | 2 | > 1846-1852 (7aa), score: 0.717<br>LTPNGLE |
| | 3 | > 1862-1907 (46aa), score: 0.695<br>KRWGRAINYTGWLTLDGNKYYFQSNSKAVTGLQKISDKYYYFNDNG |
| | 4 | >1829-1839 (11aa), score: 0.688<br>ERYYFNPNTGV |

**Table S5.** Conformational B-cell epitopes of *C.novyi* ATX. Highlighted epitopes show that the main epitopic region for ATX is between 1200-2178 residues. The longest epitopic regions are 1800-2000 residues.

| Fragment name | Amino Acid Sequence (5'…3') | length |
|---|---|---|
| ETXF (199-302 aa) | QDILVPANTTVEVIAYLKKVNVKGNVKLVGQVSGSEWGEIPSYLA FPRDGYKFSLSDTVNKSDLNEDGTININGKGNYSAVMGDELIVKV RNLNTNNVQEYVIP | 104 |
| ATXF (1822-1992 aa) | GYQEIEGERYYFNPNTGVQESGVFLTPNGLEYFTNKHASSKRWG RAINYTGWLTLDGNKYYFQSNSKAVTGLQKISDKYYYFNDNGQ MQIKWQIINNNKYYFDGNTGEAIIGWFNNNKERYYFDSEGRLLT GYQVIGDKSYYFSDNINGNWEEGSGVLKSGIFKTPSGFKL | 171 |
| E1 | IVKYRSLSIKAPGIK | 15 |
| E2 | YSAVMGDELIVKVRN | 15 |
| A1 | ITREQLMKIASIPLK | 15 |
| A2 | LNSAMLMQLLIDYKP | 15 |
| PADRE | AKFVAAWTLKAAA | 13 |
| Linker for B-cell epitopes (BL) | GGSSGG | 6 |
| Linker for Helper T-cell epitopes (TL) | GPGPG | 5 |
| TEF (A1, E1, A2, E2) | GGSSGGGPGPGITREQLMKIASIPLKGPGPGIVKYRSLSIKAPGIK GPGPGLNSAMLMQLLIDYKPGPGPGYSAVMGDELIVKVRNGPG PGGGSSGG | 97 |

**Table S6.** Different selected protein fragments from *C. perfringens* ETX and *C. novyi* ATX to design a recombinant fusion protein as a vaccine candidate against both bacteria. ATXF: Selected fragment from *C. novyi* ATX. A1, A2: The best-scored Helper T-cell epitopes for ATX. ETFX: Selected fragment from *C. perfringens* ETX. E1, E2: The best-scored Helper T-cell epitopes for ETX.

| **Model** | **Fusion Protein (FP) Design** |
|---|---|
| rFPA | PADRE + TEF + ATXF + ETXF |
| rFPB | PADRE+ ATXF + ETXF + TEF |
| rFPC | PADRE + A1 + A2 + ATXF + ETXF + E1 + E2 |
| rFPD | PADRE + ATXF + TEF + ETXF |
| rFPE | PADRE + ETXF + TEF + ATXF |

**Table S7.** Different designs for a recombinant fusion protein from epitopic fragments and MHC-II binding epitopes of *C. perfringens* ETX and *C. novyi* ATX. PADRE: Adjuant sequence. TEF: T-helper Epitopic Fragment including A1, E1, A2, E2. ATXF: Selected fragment from *C. novyi* ATX. A1, A2: The best-scored Helper T-cell epitopes for ATX. ETFX: Selected fragment from *C. perfringens* ETX. E1, E2: The best-scored Helper T-cell epitopes for ETX.

| Selected Model | Amino Acid Sequence (5'…3') | Length |
|---|---|---|
| rFPD<br><br>[Methionine<br>+ PADRE<br>+ ATXF<br>+ TEF (A1, E1, A2, E2)<br>+ ETXF] | MAKFVAAWTLKAAAGYQEIEGERYYFNPNTGVQESGVFLTPNG<br>LEYFTNKHASSKRWGRAINYTGWLTLDGNKYYFQSNSKAVTGLQ<br>KISDKYYYFNDNGQMQIKWQIINNNKYYFDGNTGEAIIGWFNN<br>NKERYYFDSEGRLLTGYQVIGDKSYYFSDNINGNWEEGSGVLKSG<br>IFKTPSGFKLGGSSGGGPGPGITREQLMKIASIPLKGPGPGIVKYRS<br>LSIKAPGIKGPGPGLNSAMLMQLLIDYKPGPGPGYSAVMGDELIV<br>KVRNGPGPGGGSSGGQDILVPANTTVEVIAYLKKVNVKGNVKLV<br>GQVSGSEWGEIPSYLAFPRDGYKFSLSDTVNKSDLNEDGTINING<br>KGNYSAVMGDELIVKVRNLNTNNVQEYVIP | 386 aa |

**Table S8.** The protein sequence of rFPD consists of 386 amino acid residues and starts with methionine attached to the N-terminal end, serving as the initial amino acid. The adjuvant, PADRE sequence, is highlighted in red for clarity. TEF, representing T-helper Epitopic Fragment, encompasses regions A1, E1, A2, and E2, crucial for eliciting immune response. ATXF, denoting selected fragments from *C. novyi* ATX, is depicted in brown, along with A1 and A2, the highest-scoring Helper T-cell epitopes for ATX. Similarly, ETFX, selected fragments from *C. perfringens* ETX, and E1 and E2, the top-scoring Helper T-cell epitopes for ETX, are depicted in green. GGSSGG serves as the linker for B-cell epitopes, shown in purple, while the linker for Helper T-cell epitopes contains the GPGPG sequence, represented in blue.

| Name | Amino Acid Sequence (5'…3') | Length (bp) |
|---|---|---|
| ETXF<br><br>(199-302 aa) | caagatatactagtaccagctaatactactgtagaagtaatagcatatttaaaaa<br>aagttaatgttaaaggaaatgtaaagttagtaggacaagtaagtggaagtgaatg<br>gggagagatacctagttatttagcttttcctagggatggttataaatttagtttatcg<br>gatacagtaaataagagtgatttaaatgaagatggtactattaatattaatggaa<br>aaggaaattatagtgcagttatgggagatgagttaatagttaaggttagaaattta<br>aatacaaataatgtacaagaatatgtaatacct | 312 |
| ATXF<br><br>(1822-1992 aa) | ggatatcaagagatagagggagaaaggtattattttaatcctaatactggagttca<br>agaatcaggagtgtttcttacgccaaatggactagaatattttacaaataaacatg<br>caagctccaaaagatgggggcgagctataaattatactggttggttgactttggat<br>ggaaataaatactattttcaatctaatagtaaagcagtaacaggattacaaaaaa<br>tatctgataaatattattactttaatgataatggacaaatgcaaataaaatggcaa<br>attataaataacaataaatattattttgatggaaatacaggcgaagctataattgg<br>atggtttaataataataaagaaagatattattttgatagtgaaggtagacttttaac<br>aggatatcaagttataggagataaatcatattatttttcagataatataaatggaa<br>attgggaagaaggaagtggagtgttaaaaagtggtattttcaaaactccttctgga<br>tttaaactt | 513 |
| E1 | atagtaaaatataggagtctttctattaaggcaccaggaataaaa | 45 |
| E2 | tatagtgcagttatgggagatgagttaatagttaaggttagaaat | 45 |
| A1 | ataacaagagaacaattaatgaaaattgcaagtataccattaaaa | 45 |
| A2 | ttaaattcagctatgttaatgcaattattaatagattataagcct | 45 |
| PADRE | gccaagttcgtggccgcctggaccctgaaggccgccgcc | 39 |

| | | |
|---|---|---|
| Linker for B-cell epitopes (BL) | ggcggcagcagcggcggc | 18 |
| Linker for Helper T-cell epitopes (TL) | ggccccggccccggc | 15 |
| TEF | ggcggcagcagcggcggcggccccggccccggcataacaagagaacaattaat<br>gaaaattgcaagtataccattaaaaggccccggccccggcatagtaaaatatagg<br>agtctttctattaaggcaccaggaataaaaggccccggccccggcttaaattcagc<br>tatgttaatgcaattattaatagattataagcctggccccggccccggctatagtgc<br>agttatgggagatgagttaatagttaaggttagaaatggccccggccccggcggc<br>ggcagcagcggcggc | 291 |
| rfpd<br><br>[atg<br>+ PADRE<br>+ ATXF<br>+ TEF (A1, E1, A2, E2)<br>+ ETXF<br>+ tag] | atggccaagttcgtggccgcctggaccctgaaggccgccgccggatatcaagaga<br>tagagggagaaaggtattattttaatcctaatactggagttcaagaatcaggagtg<br>tttcttacgccaaatggactagaatattttacaaataaacatgcaagctccaaaag<br>atgggggcgagctataaattatactggttggttgactttggatggaaataaatact<br>attttcaatctaatagtaaagcagtaacaggattacaaaaaatatctgataaatat<br>tattactttaatgataatggacaaatgcaaataaaatggcaaattataaataaca<br>ataaatattattttgatggaaatacaggcgaagctataattggatggtttaataat<br>aataaagaaagatattattttgatagtgaaggtagacttttaacaggatatcaagt<br>tataggagataaatcatattatttttcagataatataaatggaaattgggaagaag<br>gaagtggagtgttaaaaagtggtattttcaaaactccttctggatttaaacttggcg<br>gcagcagcggcggcggccccggccccggcataacaagagaacaattaatgaaa<br>attgcaagtataccattaaaaggccccggccccggcatagtaaaatataggagtc<br>tttctattaaggcaccaggaataaaaggccccggccccggcttaaattcagctatg<br>ttaatgcaattattaatagattataagcctggccccggccccggctatagtgcagtt<br>atgggagatgagttaatagttaaggttagaaatggccccggccccggcggcggca<br>gcagcggcggccaagatatactagtaccagctaatactactgtagaagtaatagc<br>atatttaaaaaaagttaatgttaaaggaaatgtaaagttagtaggacaagtaagt<br>ggaagtgaatggggagagatacctagttatttagcttttcctagggatggttataa<br>atttagtttatcggatacagtaaataagagtgatttaaatgaagatggtactatta<br>atattaatggaaaaggaaattatagtgcagttatgggagatgagttaatagttaa<br>ggttagaaatttaaatacaaataatgtacaagaatatgtaataccttag | 1161 |

**Table S9.** This table presents the reverse translation of the Recombinant Fusion Protein D (rFPD) into the corresponding nucleotide sequence, referred to as the *rfpd* gene. The nucleotide sequence of the *rfpd* gene comprises 1161 base pairs (bp). The initiation codon "ATG" signifies the start codon located at the N-terminal end, initiating the translation process, while the termination codon "TAG" denotes the stop codon positioned at the C-terminal end, signifying the end of protein synthesis. Highlighted in red, the adjuvant PADRE sequence is prominently displayed for clarity, indicating its pivotal role in enhancing immunogenicity. Additionally, the table delineates specific regions within the *rfpd* gene, such as the T-helper Epitopic Fragment (TEF), comprising regions A1, E1, A2, and E2, crucial for eliciting an immune response. Moreover, the table categorizes selected fragments from *C. novyi* ATX, as ATXF, depicted in brown, along with the highest-scoring Helper T-cell epitopes (A1 and A2) for ATX. Similarly, ETFX, selected fragments from *C. perfringens* ETX, denoted as ETFX, are represented in green, along with the top-scoring Helper T-cell epitopes (E1 and E2) for ETX. The nucleotide sequence of the linkers for B-cell epitopes depicted in purple, facilitating the structural integrity and functionality of the fusion protein. The sequence for the linkers of Helper T-cell epitopes, is represented in blue, ensuring proper spatial arrangement and interaction with immune cells.
